## Supplemental Figures 1-5 for "Mutant p53 Drives Clonal Hematopoiesis through Modulating Epigenetic Pathway"

**Supplementary Information**

**SUPPLEMENTARY FIGURE LEGENDS**

**Supplementary Figure 1.** (**a**) Bone marrow (BM) cell numbers from *p53^+/+^*, *p53^+/-^*, *p53^-/-^*, and *p53^R248W/+^* mice (8 to 12 weeks old). n=6-7 mice per group. (**b**) The absolute number of LSKs in two femurs of *p53^+/+^*, *p53^+/-^*, *p53^-/-^*, and *p53^R248W/+^* mice. n=6-7 mice per group. (**c**) The absolute number of LT-HSCs (CD48^-^CD150^+^LSKs) in two femurs of *p53^+/+^*, *p53^+/-^*, *p53^-/-^*, and *p53^R248W/+^* mice. n=6-7 mice per group. (**d**) Lineage contribution of donor-derived cells in the peripheral blood (PB) of the primary recipient mice at 20 weeks following HSC transplantation was determined by flow cytometry analysis. n =7 mice per group. (**e**) Lineage contribution of donor-derived cells in the BM of the primary recipient mice at 20 weeks following HSC transplantation was determined by flow cytometry analysis. n =7 mice per group. (**f**) Percentage of donor-derived cells in the PB of recipient mice following secondary transplantation. n=7 mice per group. (**g**) Percentage of donor-derived cells in the BM of recipient mice at 20-week after secondary transplantation. n=7 mice per group. (**h**) Lineage contribution of donor-derived cells in the PB of recipient mice at 20 weeks following secondary transplantation was determined by flow cytometry analysis. n =7 mice per group. (**i**) 1 x 10^7^ BM cells (CD45.2^+^) from *p53*^+/+^ and *p53^R248W/+^* mice were transplanted into lethally irradiated recipient mice (CD45.1^+^). At 18 hours after transplantation, the percentage of donor-derived cells in the BM of recipient mice was determined by flow cytometry. n=5-6 mice per group. (**j**) The frequency of donor-derived Lin^-^Sca1^+^CD48^-^CD150^+^ cells in the BM of recipient mice 18 hours after transplantation was determined by flow cytometry. n=5-6 mice per group. Data are represented as mean ± SEM. *P*-values calculated by ordinary one-way ANOVA test. **P*<0.05, ***P*<0.01, ****P*<0.001, *****P*<0.0001.

**Supplementary Figure 2.** (**a**) Kaplan-Meier survival curve of *p53*^+/+^ and *p53^R248W/+^* mice after 9Gy total body irradiation (TBI). n = 10 mice per group. (**b**) Fluorescence intensity of γ-H2AX in LT-HSCs at 2 hours after 2Gy irradiation was determined by ImageStream flow cytometry analysis. (**c**) γ-H2AX foci generation in HSCs following irradiation. LT-HSCs from *p53*^+/+^ and *p53^R248W/+^* mice were immunostained for γ-H2AX at 2 hours after 2Gy irradiation. LT-HSCs were stained with DAPI to identify the nuclei. (**d**) The frequency of donor-derived MEPs, CMPs, and GMPs in the BM of primary recipient mice 16 weeks following transplantation was determined by flow cytometry analysis. n = 7 mice per group. (**e**) Lineage contribution of donor-derived cells in the PB of the primary recipient mice 16 weeks following transplantation was determined by flow cytometry analysis. n = 7 mice per group. Data are represented as mean ± SEM. *P*-values calculated by ordinary one-way ANOVA test. ****P*<0.001.

**Supplementary Figure 3.** (**a**) Gene Set Enrichment Analysis (GSEA) identified hematopoietic stem cell (HSC) related gene sets are enriched in p53 mutant HSPCs compared to wild-type HSPCs in Molecular Signature Database (MsigDB). (**b**) GSEA analysis show enrichment of AML signatures in p53 mutant HSPCs compared to wild-type HSPCs. (**c**) DAVID pathway analysis of genes upregulated in p53 mutant HSPCs compared to wild-type HSPCs*.* (**d**) Quantitative RT-PCR analysis of the mRNA levels of *MLL1, MLL2,* and *MOZ* in HSCs. n=3 biological replicates. (**e**) GREAT pathway analysis of non-overlapping H3K27me3 peaks in *p53*^R248W/+^ HSPCs. (**f**) GREAT pathway analysis of non-overlapping H3K27me3 peaks in *p53*^+/+^ HSPCs. (**g**) Quantitative RT-PCR analysis of mRNA levels of *Gadd45g* in HSPCs with or without TPO stimulation. n=3 biological replicates. (**h**) Ectopic *Gadd45g* expression decreases the colony formation of p53 mutant BM cells. n = 3 independent experiments performed in duplicate. Data are represented as mean ± SEM. *P*-values calculated by ordinary one-way ANOVA test. **P*<0.05, ****P*<0.001.

**Supplementary Figure 4.** (**a**) The expression of the components of the PRC2 complex, including *Ezh1*, *Ezh2*, *Eed*, and *Suz12*, in HSPCs was determined by quantitative RT-PCR analysis. n=3 biological replicates. (**b**) Ectopic expression of wild-type (WT) or mutant p53 did not affect the protein levels of the PRC2 components in 32D cells. (**c**) Median fluorescent intensity (MFI) of p53 in the nucleus of *p53*^+/+^ and *p53*^R248W/+^ HSPCs. n=3 biological replicates. (**d**) MFI of Ezh2 in the nucleus of *p53*^+/+^ and *p53*^R248W/+^ HSPCs. n=3 biological replicates. Data are represented as mean ± SEM. *P*-values calculated by ordinary one-way ANOVA test. ***P*<0.01.

**Supplementary Figure 5.** (**a**) *Gadd45g* expression in *p53*^+/+^, *Ezh2^+/-^*, *p53*^R248W/+^ and *p53*^R248W/+^ *Ezh2^+/-^* HSPCs. n=3 biological replicates. (**b**) The absolute number of donor-derived CMPs in one femur and one tibia of recipient mice 20 weeks following pI:pC treatment. n =6-7 mice per group. (**c**) The absolute number of donor-derived MEPs in one femur and one tibia of recipient mice 20 weeks following pI:pC treatment. n =6-7 mice per group. (**d**) The absolute number of donor-derived GMPs in one femur and one tibia of recipient mice 20 weeks following pI:pC treatment. n =6-7 mice per group. Data are represented as mean ± SEM. *P*-values calculated by ordinary one-way ANOVA test. ***P*<0.01, ****P*<0.001, *****P*<0.0001.

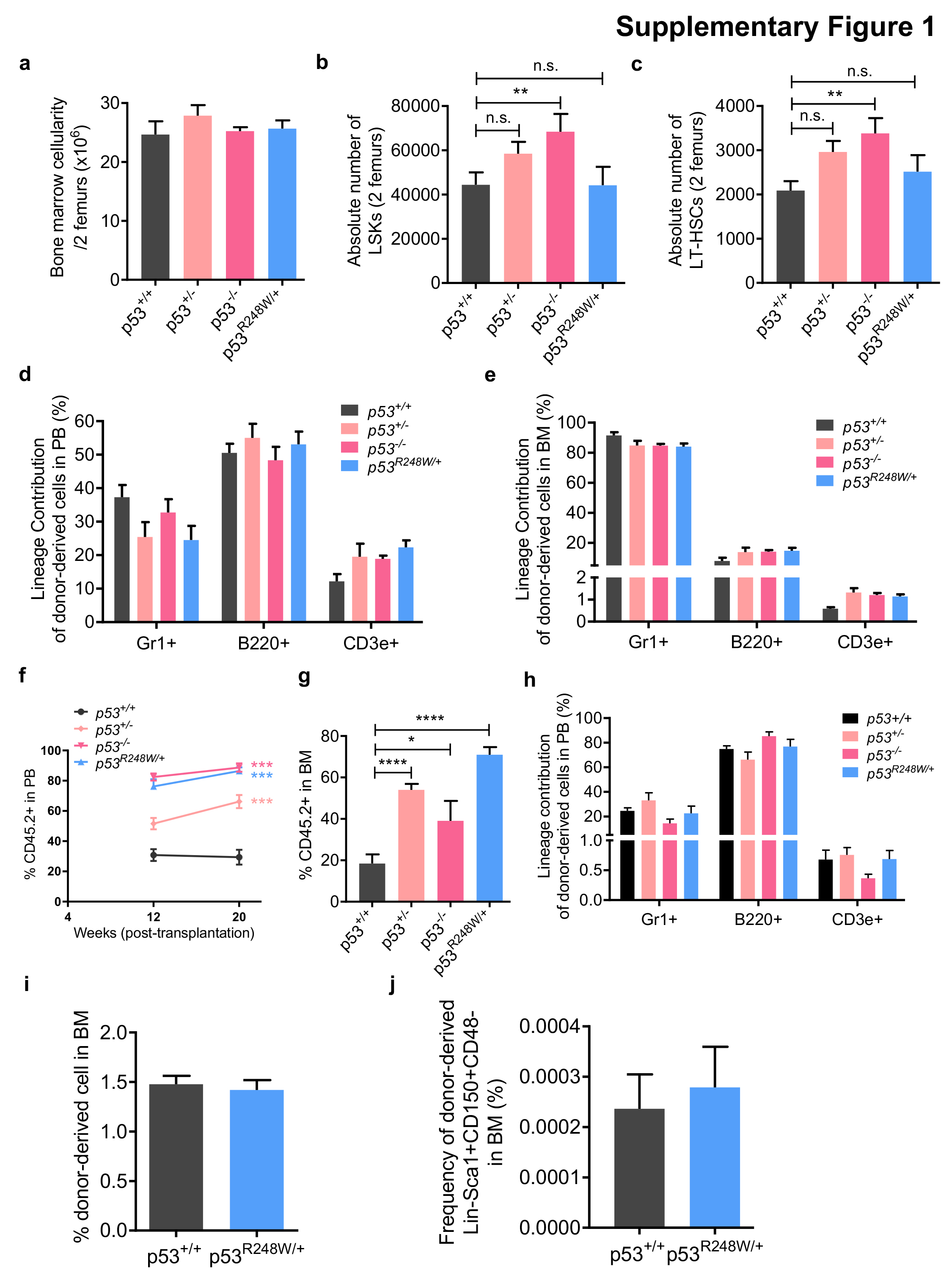

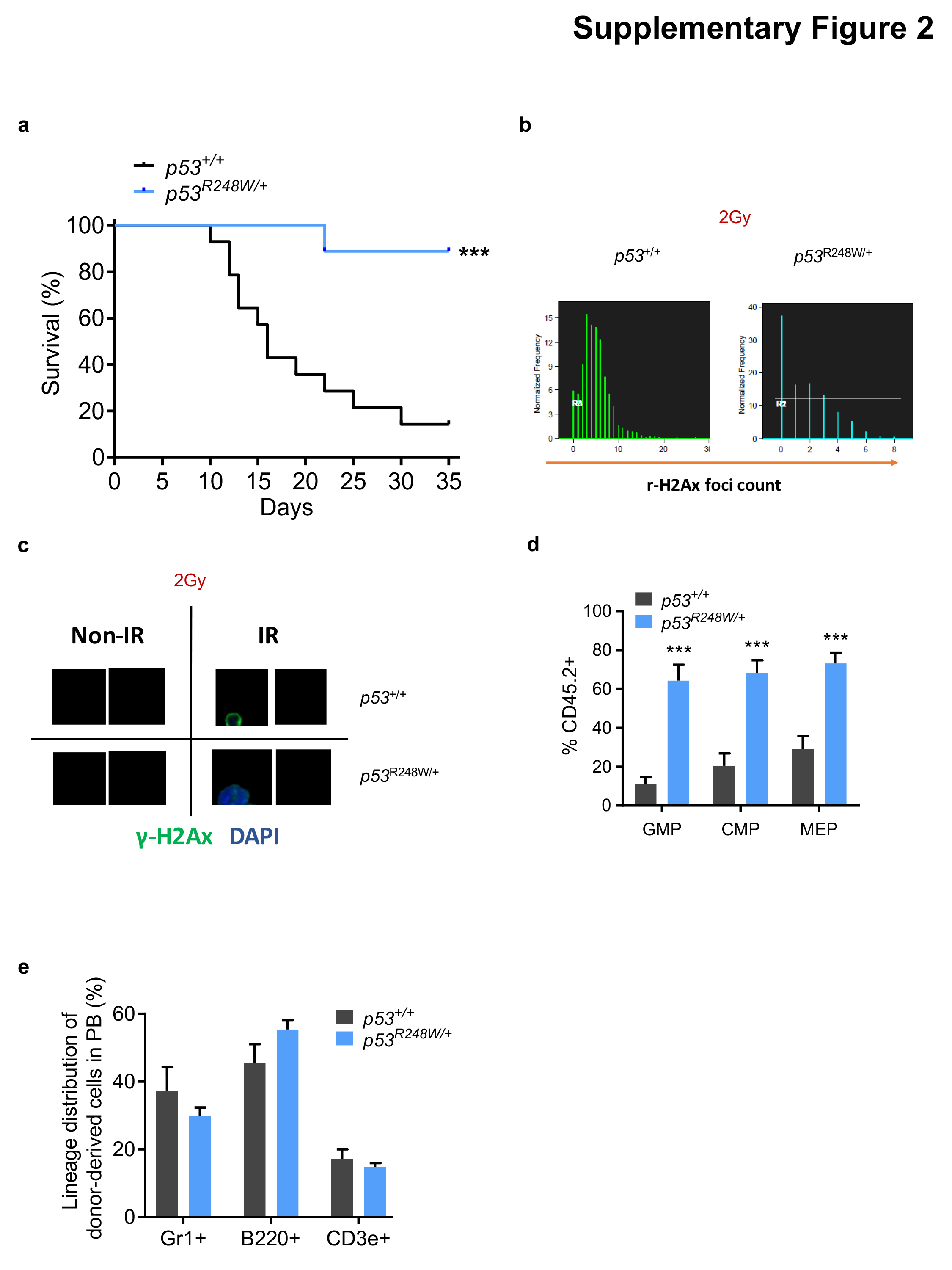

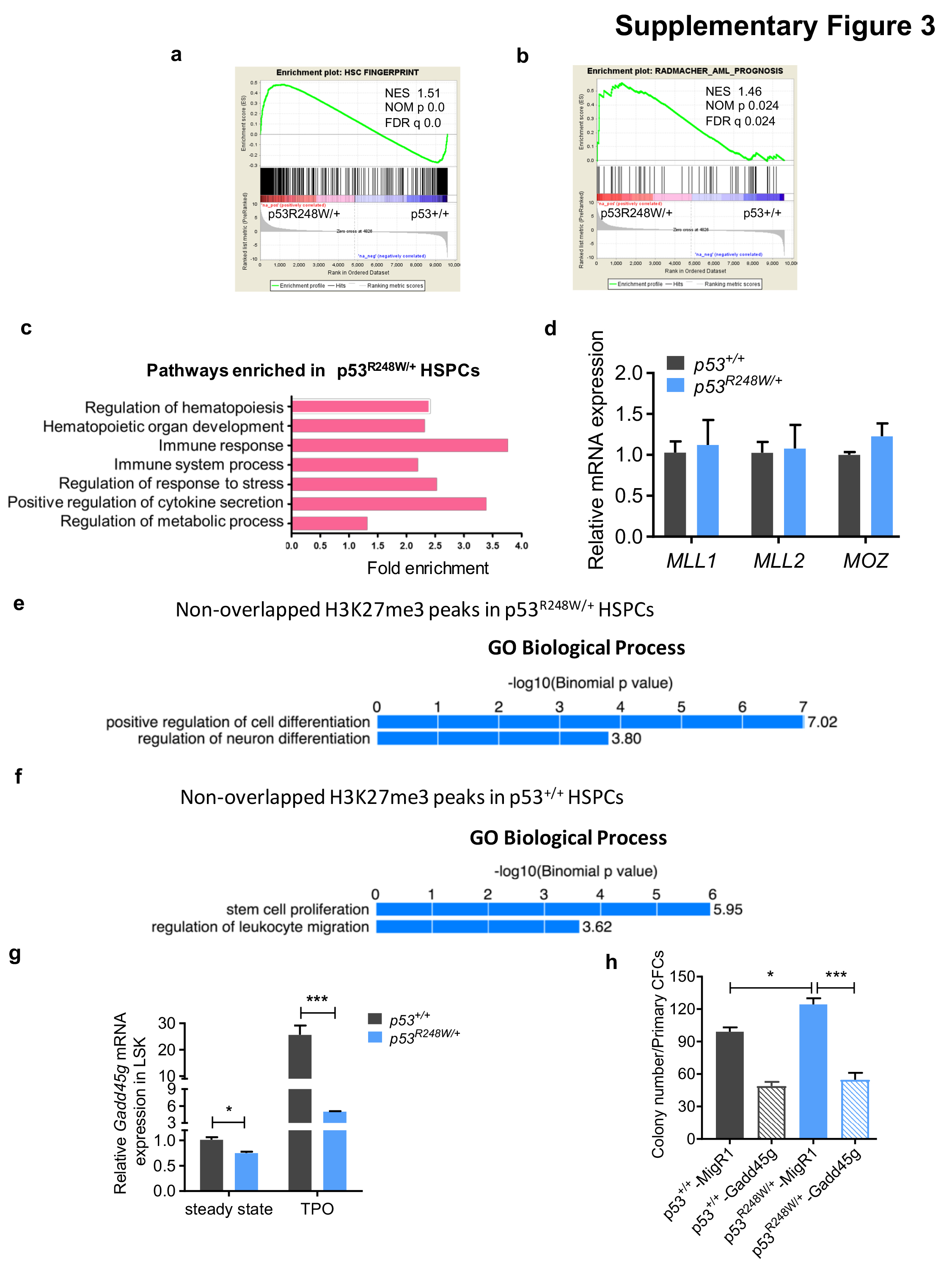

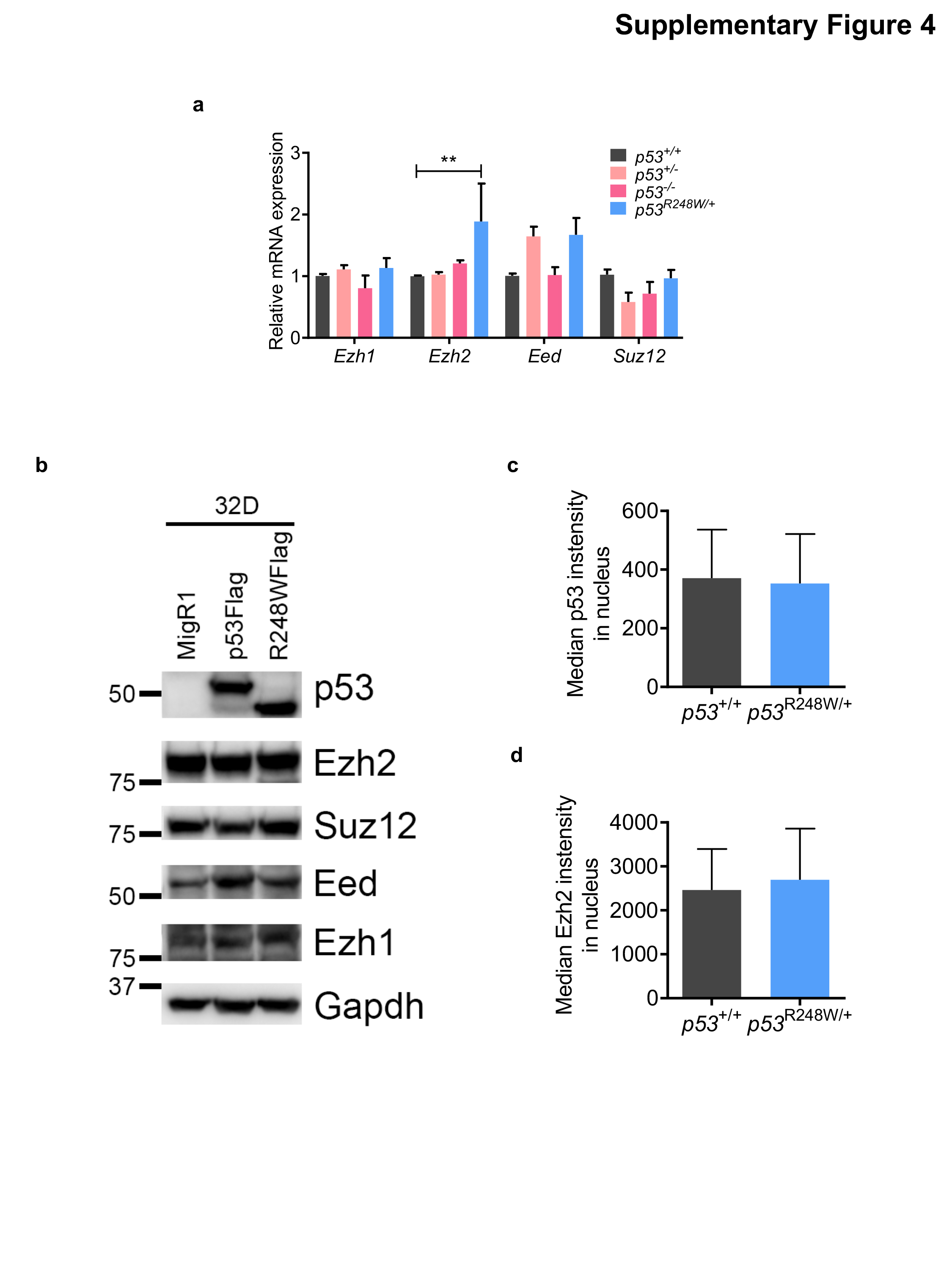

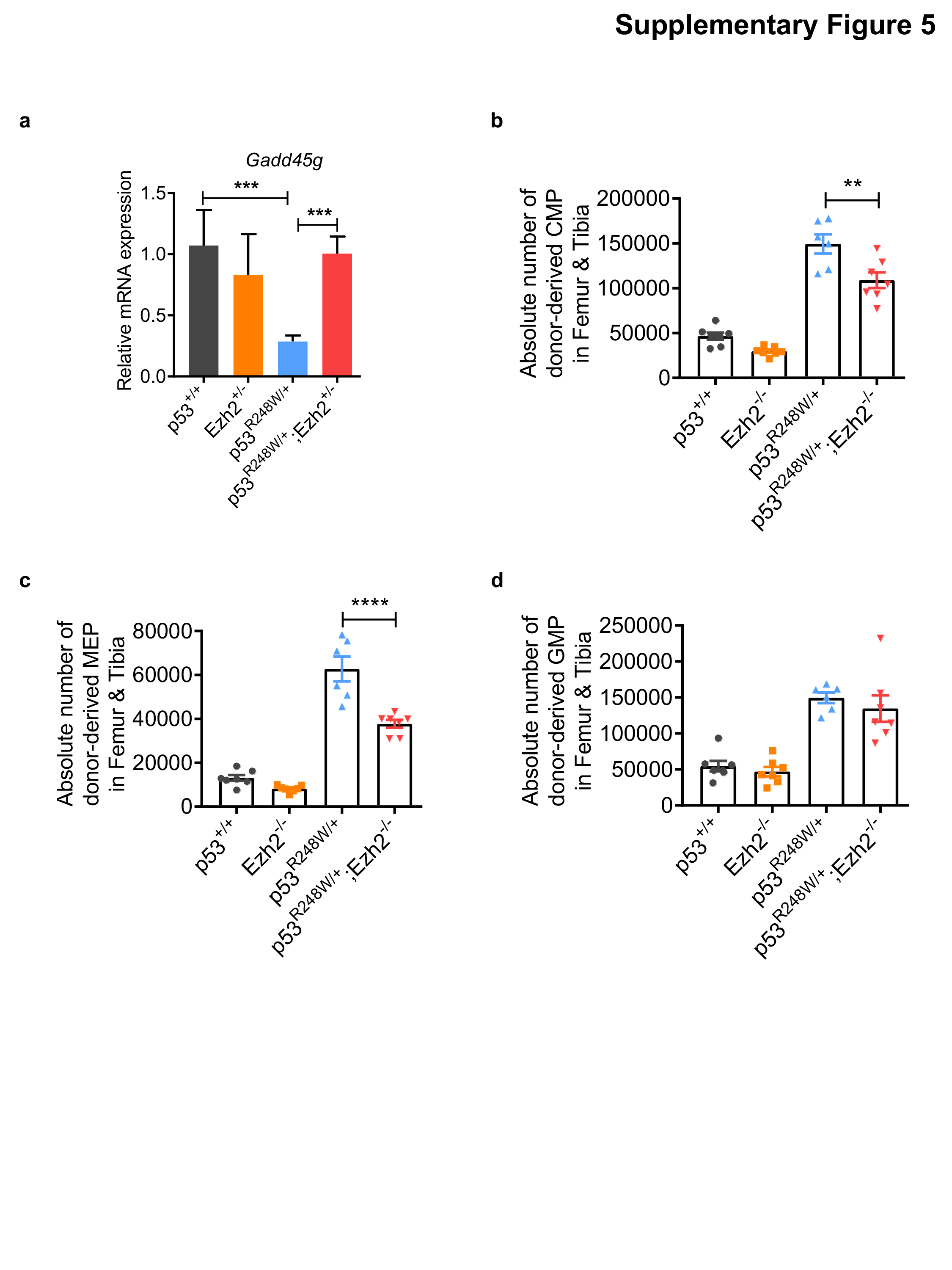
